# Functional properties of powders produced from either or not fermented mealworm (*Tenebrio molitor*) paste

**DOI:** 10.1101/2020.04.17.042556

**Authors:** A. Borremans, S. Bußler, T. S Sagu, H.M. Rawel, O. Schlüter, L. Van Campenhout

## Abstract

The aim of this study was to determine the effect of fermentation of mealworms (*Tenebrio molitor*) with commercial meat starters cultures on the functional properties of powders produced from the larvae. Full fat and defatted powder samples were prepared from non-fermented and fermented mealworm pastes. Then the crude protein, crude fat and dry matter contents, pH, bulk density, colour, water and oil binding capacity, foaming capacity and stability, emulsion capacity and stability, protein solubility, quantity of free amino groups and protein composition of the powders were evaluated. Regardless of the starter culture used, fermentation significantly (p < 0.05) reduced the crude and soluble protein content of the non-defatted mealworm powders and in general impaired their water and oil binding, foaming- and emulsifying properties. Defatting of the powders improved most functional properties studied, except the protein solubility, water binding capacity, foaming capacity and emulsion stability. The o-phthaldialdehyd assay revealed that the amount of free amino groups increased during fermentation, which may be attributed to proteolysis of mealworm proteins by the starters. Sodium dodecyl sulfate polyacrylamide gel electrophoresis demonstrated that the soluble proteins of fermented powders were composed of molecules of lower molecular mass compared to non-fermented powders. As the molecular sizes of the soluble proteins decreased, it is clear that also the protein structure was modified by the fermentation process, which in turn led to changes in functional properties. It was concluded that fermentation of mealworms in general does not contribute to the functional properties studied in this work. Nevertheless, the results confirmed that the properties of non-fermented powders are comparable to other food protein sources.

## 1. Introduction

In recent years, there is an increased interest in the use of edible insects for food applications. In particular, mealworms (*Tenebrio molitor*) are gaining attention as alternative protein source due their high protein level, good amino acid profiles and high levels of unsaturated fatty acids, vitamins and minerals (Van Huis, 2016). Various technologies are used for the stabilisation of mealworms and processing them into foods, most of which are based on heat application (oven drying and boiling). It can be interesting, both cost-wise and protein property-wise, to apply processes that do not involve heat. Fermentation is a non-thermal process in which a food matrix is subjected to the action of microorganisms or enzymes so that desirable biochemical changes cause modification to the product (Campbell-Platt, 1987). These modifications may result in modified sensory qualities, improved nutritional value, enhanced preservation and/or increased economic value. Fermentation of mealworm paste has been reported to be feasible with lactic acid starter cultures, as indicated by a rapid pH reduction (De Smet et al., 2019; Borremans et al., 2019). During fermentation, some of the starter cultures tested generated free glutamic and aspartic acid, two amino acids that are responsible for the well-appreciated umami taste. The impact of fermentation on other properties of the paste was not investigated so far and is the subject of this study. Fermentation is expected to alter the characteristics of the insect proteins, but it is not known how this translates into their nutritional value and technological functionality and hence in their application potential as food ingredients.

In Western countries, insects are believed to be better accepted by consumers when they are fragmented and included in a food as an ingredient, rather than in their whole form. For mealworms to be successful in food applications, they should ideally possess several desirable characteristics, referred to as functional properties. To date, only a few studies have considered the technical functionality of mealworms or protein preparations thereof, such as solubility, water and oil binding, gelling, foaming and emulsifying capacity (Bußler et al., 2016; Kim et al., 2016; Yi et al., 2013; Xue Zhao et al., 2016; Zielińska et al., 2018). In general, mealworms were found to have high water and oil binding, good emulsion properties, but moderate to poor foam and gelling properties. Their proteins exhibited good solubility in the pH range of 2 to 12, making them a suitable candidate for many food applications. Despite several studies on the functional properties of mealworms, the effects of fermentation on these properties are unclear. Some properties may be affected by fermentation as fermentation tends to alter the structure of proteins, which are the main functional constituents in emulsions, foams and gels (Broyard & Gaucheron, 2015).

The objective of this study was to investigate the impact of fermentation on the functional properties of mealworm powders. Mealworm pastes were produced and, apart from the control samples, either fermented with the commercial meat starter culture Bactoferm^®^ F-LC or with the pure culture *Lactobacillus farciminis*. Both starters have the generally-recognised-as-safe (GRAS) status and showed promising results in previous research (i.e. rapid acidification and inhibition of undesirable microorganisms). Full fat and defatted powders were produced from all pastes and characterised with respect to moisture content, proximate composition, pH, bulk density, and colour. To evaluate the fermentation-induced effects on protein functionality, water and oil binding, foaming and emulsion properties as well as protein solubility (pH 2 to 10) of protein extracts recovered from defatted and non-defatted powders were analysed. Underlying mechanisms for the fermentation effects observed were studied by analysing free amino groups and the molecular weight distribution of (soluble) proteins.

## 2. Materials and methods

### 2.1 Sample preparation and processing

Mealworms (*Tenebrio molitor* larvae) were purchased from the commercial supplier Nusect (Sint-Eloois-Winkel, Belgium). The living mealworms were packaged in freezer bags (3 x 1.2 kg mealworms/bag), killed by freezing and stored at −18 °C until further use.

Mealworm powders (non-fermented, fermented and defatted samples) were prepared as shown in **Figure 1A**. Non-fermented samples were prepared by thawing one bag of mealworms for 4 hours at 4°C and mixing the larvae into a paste using a kitchen mixer as described earlier (Borremans et al., 2019). The larvae of the other two bags were blanched, mixed into a paste, triplicate volumes of 50 g were inoculated with either the commercial meat starter culture Bactoferm^®^ F-LC (Chr. Hansen, Denmark, mixture of *Staphylococcus xylosus, Lactobacillus curvatus* and *Pediococcus acidilactici*) or the pure culture *Lactobacillus farciminis* (kindly provided by Chr. Hansen), and subsequently fermented according to Borremans et al. (2018, 2019). Powders were produced by freeze drying (48 h, Büchi Lyovapor L-200) and grinding (60s, Clatronic KSW 3307). To prepare defatted powders, solvent extraction using n-hexane was performed. A proportion of each powder was mixed with hexane (1:10 ratio, v/w) and stirred on a magnet stirrer for 1 h. After sedimentation of the solids, the hexane-fat-mixture was decanted and residual hexane was removed by evaporation overnight.

**Figure 1.**
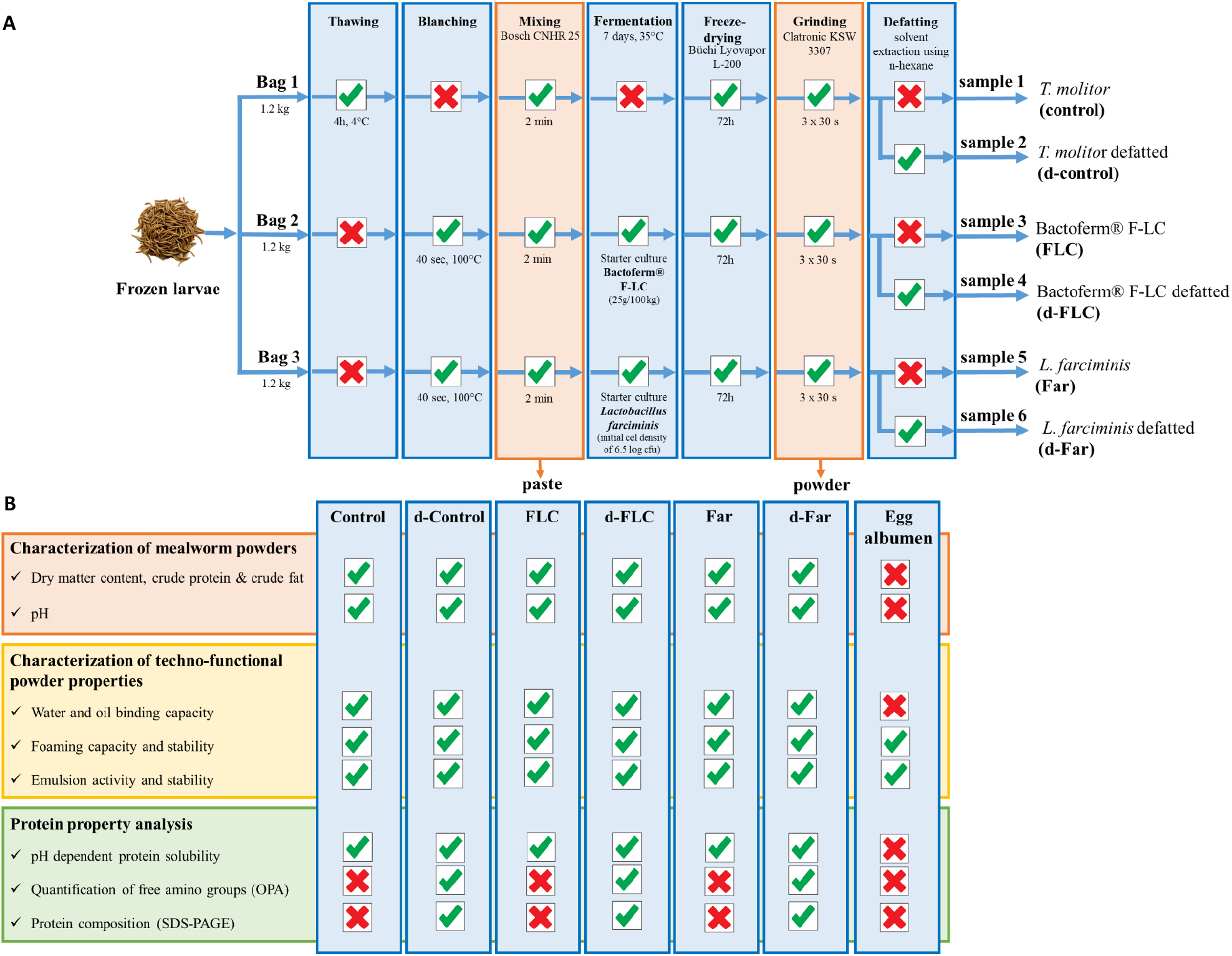
Schematic representation of sample preparation (A) and overview of the performed analyses (B).

**Figure 1B** gives an overview of the performed analyses. The water- and oil binding capacity of the powders were determined at their native pH, the foaming and emulsifying properties and the free amino groups at pH 7 (indicative of a neutralisation process) and protein solubility and protein composition were determined in the pH range of 2 to 12. Egg albumen powder served as reference material from Pulviver (Bastogne, Belgium).

### 2.2 Characterization of mealworm powders

#### 2.2.1 Crude protein, crude fat, dry matter content

Crude protein and crude fat contents were determined in triplicate as described by Bußler et al. (2016). Crude protein contents (NKjel, conversion factor 6.25) were determined using the method by Kjeldahl (Kjeldatherm Turbosog, Titrino plus 848, Gerhardt Analytical Systems, Königswinter, Germany), according to DIN EN 25663: Digestion and distillation (Kjeldahl Sampler System K-370/371) were conducted as described by the Association of German

Agricultural Investigation and Research Institutions (VDLUFA, 1976). Crude fat content of the flour fractions was determined according to the filter bag method Am 5-04 (AOCS 2005) as an indirect method for measuring crude oil (Filterbags XT4, ANKOM Technology, New York, USA). Dry matter contents were determined by drying the mealworm powders in a forced-air oven (Memmert, UFB500) at 105°C for 17 h.

#### 2.2.2 pH, bulk density and colour measurement

To determine pH, mealworm powders (3 x 1.0 g) were mixed with 10 mL of distilled water and vortexed for 30 s. The pH of the mixtures was measured using a pHenomenal pH 1100H meter with a SenTix 82 pH electrode (VWR). Bulk density was estimated by the modified method of Wang and Kinsella (1976). A 10.0 g sample was gently packed in a 100 ml graduated cylinder by tapping ten times on a bench top from a height of 5 cm. The final volume of the sample was recorded and the bulk density was expressed as g/ml of sample. Colour measurements were performed with a colorimeter using the CIElab scale (CR-5, Konica Minolta). Five independent measurements of a* (redness), b* (yellowness) and L* (lightness) parameters were carried out for triplicate samples of each powder type. Browning indices and colour difference (ΔE) were calculated according to the formula described by Lenaerts, Van Der Borght, Callens, & Van Campenhout (2018), using non-fermented powder as reference.

### 2.3 Analysis of the techno-functional properties of the powders

#### 2.3.1 Water and oil binding capacity

Water (WBC) and oil (OBC) binding capacity of the powders were determined in triplicate according to the method of Bußler et al. (2016) with slight modifications. In brief, 0.5 g of each powder was weighted into a pre-weighted centrifuge beaker to which 3.0 mL of distilled water or 3.0 mL of commercial rape seed oil was added. The mixtures were stirred (60 s or two times 60 s for WBC and OBC, respectively) using a propeller stirrer and an overhead agitator (Yellowline^®^, IKA^®^ OST basic, USA), and centrifuged at 3900 g for 20 min. The samples were re-weighted after discarding the supernatant and putting the beakers upside-down on absorbent paper for 60 min. The differences in weight were calculated and the results were presented as gram of water or oil absorbed per gram of powder.

#### 2.3.2 Foaming capacity and stability

For foaming capacity and stability determinations, 0.25% w/v protein suspensions were prepared at pH 7. Briefly, each sample (quantity depending on solubility of the proteins) was suspended in 100 mL distilled water and the pH was adjusted to 7 with 1.0 M NaOH or 1.0 M HCl. The solutions were stirred for 30 min at room temperature and centrifuged for 20 min (4°C, 10000 g). The supernatants were collected and stored at 4°C until subsequent analyses. The foaming properties were determined in fivefold by the method described by Zielińska et al. (2018), with modifications. Twenty milliliter of supernatant was transferred into a 250 ml beaker and each sample was individually beaten in a high shear homogenizer mixer (16000 rpm, 2 min, Ultra turrax, IKA, Staufen, Germany). The whipped sample was immediately transferred into a graduated cylinder and the total volume was read at time zero and 30 min after whipping. Foaming capacity (FC) and foaming stability (FS) were calculated using the formulas described by Zielińska et al. (2018).

#### 2.3.3 Emulsion capacity and stability

Emulsifying properties were determined in fivefold with the method of Zielińska et al. (2018), with modifications. Protein solutions (0.25% w/v, pH 7) were prepared as described earlier and 10 ml of each solution was mixed in 50 ml tubes with an equal volume of rapeseed oil dyed with liquid natural carotene (M = 536.89 g/mol, Carl Roth, Karlsruhe, Germany). Following homogenization (20000 rpm, 1 min, Ultra turrax, IKA, Staufen, Germany), the mixtures were centrifuged at 3000 g for 5 min and the volume of the individual layers were read. Emulsion stability was evaluated by heating the emulsion for 30 min at 80°C. Then, the samples were centrifuged at 3000 g for 5 min before the volume of the individual layers were read. Emulsion capacity (EC) and emulsion stability (ES) were calculated using the formulas described by Zielińska et al. (2018).

### 2.4 Analysis of the protein properties of the powders

#### 2.4.1 pH dependent protein solubility

Protein solubility was determined in the pH range of 2 to 12. Briefly, 0.1 g of each powder was mixed with 10 ml of distilled water and the pH of the mixture was adjusted to a value in the range of 2 to 12 using 1.0 M HCl or 1.0 M NaOH. The solutions were stirred on a rotary shaker (350 rpm) for 30 min and centrifuged (8.000xg, 20 min, 4°C). The protein concentration of the supernatant was assessed by the Bradford method (1976) using bovine serum albumin (Fluka, Buchs, Switserland) as a standard. The assay consisted of 800 μl of the protein extracts and 200 μl of Bradford reagent (Roti^®^-Quant, Carl Roth, Karlsruhe, Germany) reacting for 20 min at ambient temperature. The absorption maximum was analysed at 595 nm against a blank value (demineralized water) using an UV/Vis spectrophotometer (BioPhotometer plus, Eppendorf, Hamburg, Germany). The dissolving procedure and spectrophotometric measurements were each performed in triplicate and the protein solubility was calculated according to the formula described by Zielińska et al. (2018). The remainder of the protein extracts was frozen at −18°C for the quantification of free amino groups and the determination of the protein composition.

#### 2.4.2 Quantification of free amino groups

In order to quantify the amount of free amino groups in the defatted mealworm powders, the o-phthaldialdehyd (OPA) assay was used. To this end, 100 μl of protein extract (pH 7, thawed at room temperature) was added to 800 μl OPA solution and to 800 μl of the same solution but without OPA (blank). The mixtures were allowed to react at ambient temperature for 30 min before the absorbance was measured at 330 nm. Three replicates of each measurement were included in the experiment and the free amino groups were calculated against an L-leucine (Merck KGaA, Darmstadt, Germany) standard curve.

#### 2.4.3 Protein molecular weight distribution

Proteins of the defatted powders were analysed with sodium dodecyl sulfate polyacrylamide gel electrophoresis (SDS-PAGE) to determine their molecular weight distribution. Protein extracts (pH 2 to 12) were thawed at room temperature, sonicated (S10 Elmasonic, Elma Schmidbauer GmbH, Singen, Germany) for 5 minutes, and 500 μl of each replicate per pH-value were pooled.

Electrophoresis was performed under reducing and denaturing conditions using Invitrogen NuPAGE 12% Bis-Tris precast protein gels with 12 wells (Thermo Fisher Scientific, CA, USA). Non-fermented samples were mixed with NuPAGE™ LDS Sample Buffer (containing glycerol, 2-mercaptoethanol, SDS and Coomassie blue G250 and pH 8.4) at a ratio of 1:5, while the fermented samples were mixed at a ratio of 1:3 with the same sample buffer. As references, 10 mg of each powder from non-fermented and fermented samples were dissolved in 1500 μL samples buffer and diluted with sample buffer in a ratio of 1:2. After heating the mixtures at 100° C for 5 min, samples were cooled to room temperature and each 15 μl of the reference and non-fermented samples and 10 μl of fermented samples were loaded onto the gel. An aliquots of 5 μl of Page Ruler Plus pre-stained broad range standard containing a 9 protein ladder (protein composition in kDa of 10, 15, 25, 35, 55, 70, 100, 130 and 250; Thermo Fisher Scientific, CA, USA) was loaded as well. Table S1 (Supporting information) presents the initial protein concentration and the protein quantities loaded. The separation was carried out under constant current (30 mA/Gel) for 120 minutes using NuPAGE MES SDS Running Buffer. After separation, the gels were stained overnight at room temperature with Coomassie blue solution (in 10% acetic acid) and then destained with 10% acetic acid for 3 hours. The gels were scanned using Bio-5000 Professional VIS Gel Scanner (Provided by SERVA Electrophoresis GmbH, Heidelberg, Germany) and analyzed with Image Lab Software (Bio-Rad Laboratories Ltd., Hemel Hempstead, UK). Electrophoresis experiments were repeated two times.

### 2.5 Statistical analysis

SPSS statistics (IBM SPSS Statistics version 25, New York, USA) was used to statistically analyse the data generated. The data, reported as averages of at least three replicates, were subjected to one-way analysis of variances (ANOVA) to compare means. Next, Tukey’s post-hoc test was used to determine significant differences among samples. However, when variances were not equal, the Kruskall-wallis test with the Dunn-Bonferroni post-hoc test was performed. For all tests a significance level of 0.05 was considered.

## 3. Results and discussion

### 3.1 Characterization of mealworm powders

The proximate composition of the different mealworm powders relating to dry matter, crude protein and crude fat was determined and expressed on dry matter basis (**Table 1**). The Control had a dry matter content of 96.26%, and contained 49.68% of crude protein and 16.61% of crude fat. The protein and fat content were lower than the values reported for freeze-dried mealworms by Zhao et al. (2016; 51.5% and 32.9%, respectively), Bußler et al. (2016; 57.8% and 19.1%, respectively) and Lenaerts et al. (2018; 59.96% and 28.35%, respectively). This heterogeneity in proximate composition can be ascribed to differences in rearing and processing conditions as well as to differences in methods of analysis applied (Rumpold & Schlüter, 2013). Fermentation with the starters Bactoferm^®^ F-LC and *L. farciminis* did not significantly influence the dry matter content of the full fat powders. The crude protein content, on the other hand, significantly (p < 0.05) decreased, while (concomitantly) the crude fat content significantly (p < 0.05) increased with fermentation. Literature reports various effects of fermentation on proteins in other food matrices. Several studies (Çabuk et al., 2018; Ojokoh, Fayemi, Ocloo, & Nwokolo, 2015) observed an increase in pea and acha rice flour, respectively, while others (Klupsaite et al., 2017; Omowaye-Taiwo et al., 2015) reported a decrease in crude protein content during fermentation of lupine and a melon type, respectively. An increase in crude protein content may be attributed to loss of dry matter (mainly carbohydrates). A decrease, on the other hand, can be caused by degradation of proteins by microorganisms, thereby releasing peptides and amino acids (Nkhata et al., 2018). As these components are necessary for microbial growth and organic acids production by Lactic Acid Bacteria (Klupsaite et al., 2017), it is possible that the starters use the generated peptides and amino acids, thereby lowering the protein content of the fermented powders. Defatting of the mealworm powders (resulting in the powders d-Control, d-FLC and d-Far) contributed to a significant (p < 0.05) increase in the crude protein content of respectively 18.21%, 19.34% and 17.31%. Subsequent analysis of the fat content revealed residual fat values of less than 5.34% for all samples.

**Table 1.** Means ± standard deviations (n = 3) of dry matter (DM), crude protein (CP), crude fat content (CF) and pH of full fat (Control and the samples fermented with Bactoferm^®^ F-LC and *Lactobacillus farciminis*, FLC and Far) and defatted powders (d-Control and the fermented samples d-FLC and d-Far) produced from unfermented and fermented mealworm larvae (*Tenebrio molitor*).

| Powder | DM [g/100g as is] | CP [g/100 g DM] | CF [g/100 g DM] | pH [-] |
| --- | --- | --- | --- | --- |
| Control | 96.26 ± 0.43<sup>a</sup> | 49.68 ± 0.02<sup>a</sup> | 16.61 ± 0.23<sup>a</sup> | 6.28 ± 0.05<sup>a</sup> |
| FLC | 95.76 ± 0.13<sup>a</sup> | 42.60 ± 0.02<sup>b</sup> | 21.05 ± 0.43<sup>b</sup> | 4.60 ± 0.04<sup>b</sup> |
| Far | 96.08 ± 0.10<sup>a</sup> | 44.55 ± 0.04<sup>c</sup> | 20.20 ± 0.31<sup>c</sup> | 4.64 ± 0.04<sup>b,c</sup> |
| d-Control | 94.35 ± 0.21<sup>b</sup> | 67.89 ± 0.02<sup>d</sup> | 3.37 ± 0.10<sup>d</sup> | 6.28 ± 0.02<sup>a</sup> |
| d-FLC | 92.19 ± 0.07<sup>c</sup> | 61.94 ± 0.02<sup>e</sup> | 5.13 ± 0.21<sup>e</sup> | 4.72 ± 0.02<sup>c,d</sup> |
| d-Far | 92.21 ± 0.07<sup>c</sup> | 61.86 ± 0.04<sup>e</sup> | 3.17 ± 0.15<sup>d</sup> | 4.75 ± 0.02<sup>d</sup> |

<sup>a, b, c, d, e</sup> Mean values within a column with the same superscript are not statistically different (P > 0.05).

Results of pH, bulk density and colour parameters are summarized in **Table 1** as well. The pH of the Control powder was 6.28 and was unaffected by the defatting treatment. Fermentation significantly decreased (p < 0.05) the pH by the production of organic acids to values ranging from 4.56 to 4.68 for the full fat samples and from 4.70 to 4.77 for the defatted samples. Both hexane defatting as well as fermentation significantly increased (p < 0.05) the bulk density of all powders. The increase in bulk density (BD) by defatting may be caused by particles sticking to each other due to the presence of residual n-hexane, but the increase in BD by fermentation cannot be explained so far. According to Ogodo, Ugbogu, Onyeagba, & Okereke (2018), the BD usually decreases during fermentation due to the breakdown of complex compounds into simpler molecules by microorganisms. As to the colour characteristics of the full fat powders, the fermented ones had a higher browning index and thus darker colour than the control powder. The subsequent defatting process noticeably decreased the browning index from ± 70 to ± 30, and eliminated statistical differences in colour between the powders. ΔE, the total colour difference between the fermented samples and the control powders, was recorded in the range of 1.16 to 2.55, which is considered to be noticeable by a human observer (Mokrzycki & Tatol, 2011).

### 3.2 Impact of fermentation on the techno-functional properties of the powders

#### 3.2.1 Water and oil binding capacity

WBC and OBC of the different mealworm powders are depicted in **Figure 2**. The WBC and OBC of the non-fermented and full fat samples was 1.79 ± 0.05 and 1.51 ± 0.05 g/g and it was 1.62 ± 0.01 and 1.86 ± 0.05 g/g in non-fermented, defatted samples. Bußler et al. (2016) reported substantially lower WBC (0.8 g/g dry mass) and OBC (0.6 g/g dry mass) values for (non-fermented) non-defatted mealworm flour (*Tenebrio molitor*). Using a similar method of powder production to that used in this study, Zielińska et al. (2018) reported a WBC of 1.29 g/g and an OBC of 1.71 g/g for ground mealworms (*Tenebrio molitor*). The different origin (likely including a different rearing method and/or substrate) of the mealworms as well as the different methods of analysis used can explain the variations in the results from different studies. Compared to other protein sources, the mealworm powders tested in this study had higher WBC and OBC values than those of soy flours (130% and 84%, respectively) and comparable WBC and OBC to those of egg white flours (168% and 135%, respectively; Gravel & Doyen, 2020).

**Figure 2.**
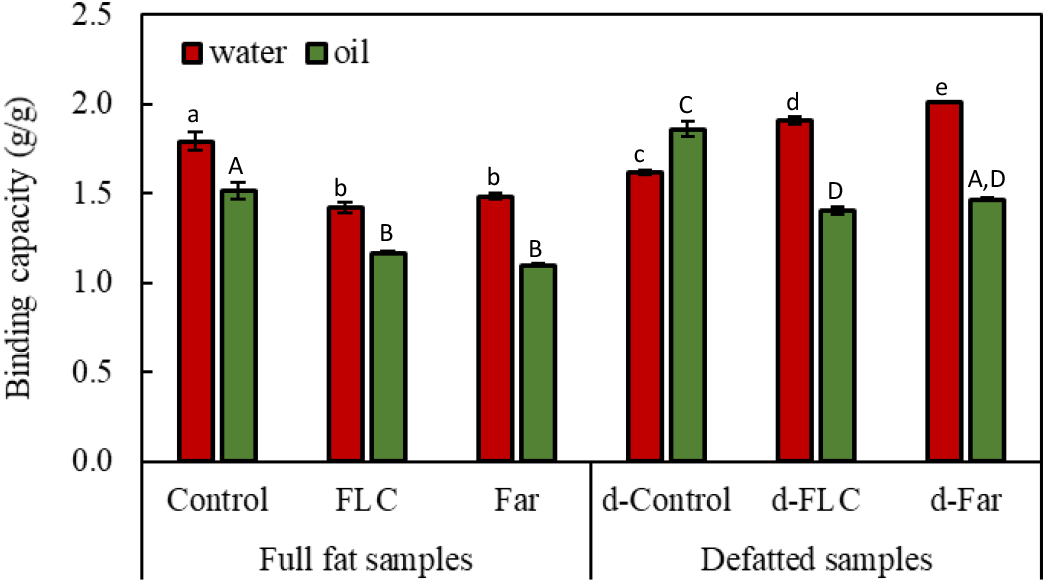
Water (WBC, red) and oil (OBC, green) binding capacity of full fat (Control, FLC and Far) and defatted (d-Control, d-FLC, d-Far) mealworm powders. Data are expressed as mean ± standard deviations (n = 3). Different letters (a,b,c,d for WBC and A,B,C,D for OBC) indicate significant (p < 0.05) differences between means.

Irrespective of the starter cultures used, fermentation induced a significant decrease in WBC (from 1.9 to 1.4-1.5 g/g) and OBC (from 1.5 to 1.1-1.2 g/g) in the full fat samples. The lower protein contents of the fermented samples compared to those of non-fermented samples (**Table 1**), as well as changes in the quality of the proteins upon fermentation, can explain this reduction in WBC and OBC. Reduction in WBC by lactic acid fermentation has been observed earlier for various flours, such as maize flour and sorghum flour (Ogodo, Ugbogu, Onyeagba, et al., 2018). Contradictory to the present study, the OBC of these flours was enhanced by fermentation with 1.2 and 0.8 mL/g, respectively. During fermentation, these proteins became partially unravelled and hence exposed buried hydrophobic groups that can bind more oil. Whereas fermentation decreased both WBC and OBC in powders that were not defatted afterwards, removing oil from the fermented mealworm powders significantly (p < 0.05) reduced OBC but increased WBC. Among all powders tested, the fermented and then defatted powders showed the highest WBC.

#### 3.2.2 Foaming capacity and stability

**Figure 3(A-B)** describes the FC and FS of the protein solutions (0.25% w/v, pH 7) extracted from the different mealworm powders. As a reference for FC and FS measurements, an egg albumen solution was used at the same concentration. Egg albumen has excellent foaming properties as it adsorbs rapidly on the air-liquid interface during whipping and rearranges to form a cohesive viscoelastic film via intermolecular interactions (Malomo et al., 2014).

**Figure 3.**
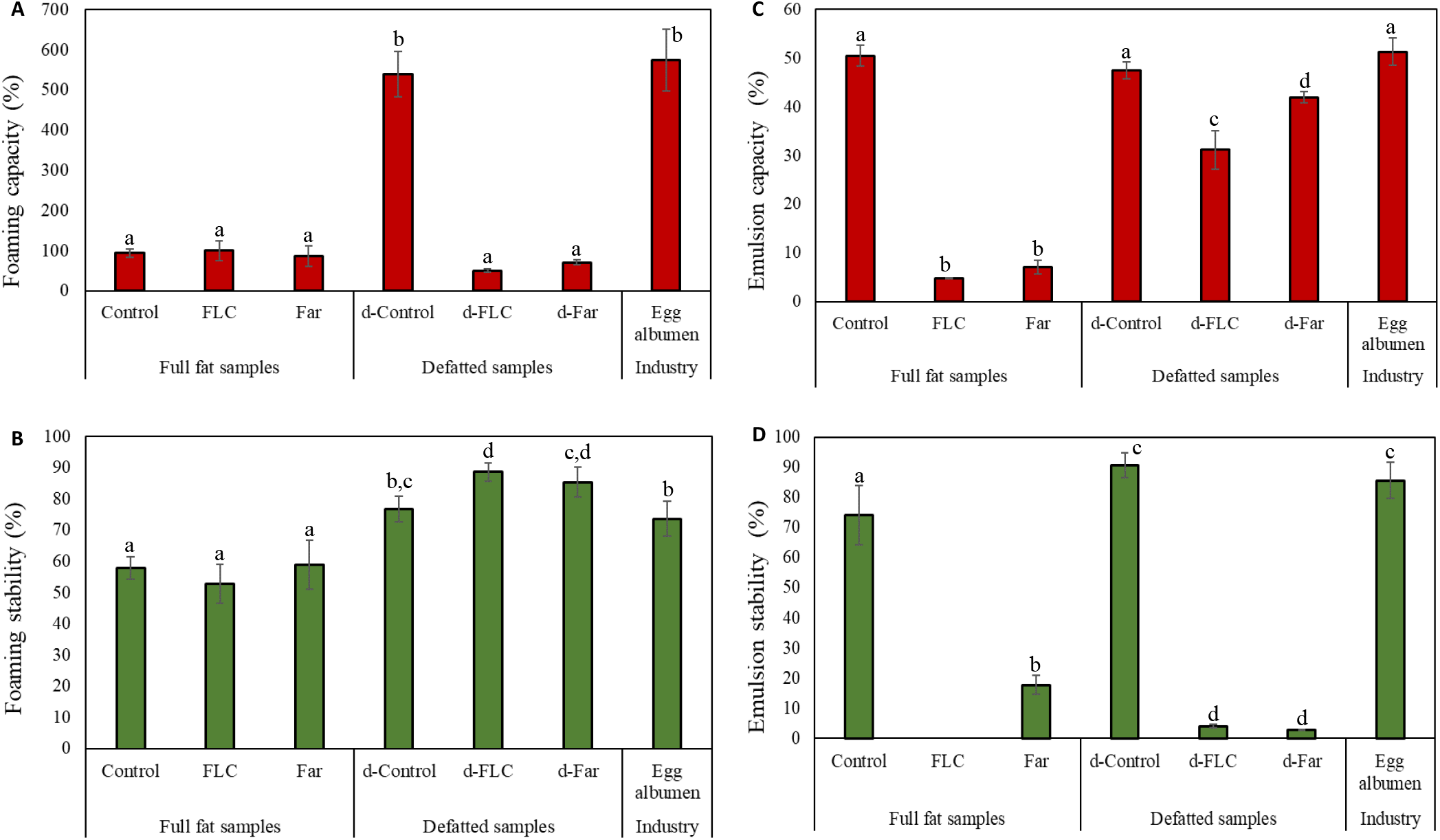
Foaming capacity (A), foaming stability (B), emulsion activity (C) and emulsion stability (D) of extracted protein solutions (0.25% w/v) from full fat (Control and the fermented samples FLC and FAR) and defatted (d-Control and the fermented samples d-FLC and d-Far) mealworm powders. Egg albumen solutions (0.25% w/v) were included as reference. Data are expressed as mean ± standard deviations (n = 5). Different letters (a,b,c,d) indicate significant (p < 0.05) differences between means.

**Figure 3A** did not reveal any significant differences in FC among the full fat powders. The FC ranged from 60 to 126%, and were low when compared with the reference (575 ± 77%). Defatting markedly improved the FC of the Control powder from 94 to 540%. This observation is consistent with the findings of Akpossan et al. (2015) that the FC of defatted flours were superior to that of full fat flours. In contrast to the Control, the FC of the fermented powders diminished upon defatting to 50 and 70%, respectively. These results demonstrate that the treatments used in this study (i.e. blanching and fermentation) may cause changes in the nature of proteins, which lead to changes in foaming properties. The FS of the powders ranged from 47 to 92% (**Figure 3B**). Similar to the FC, results of FS showed no pronounced differences among the full fat mealworm powders. The foams of the defatted powders, and especially those of the fermented powders, were significantly (p < 0.05) more stable than the foams of the full fat powders and the reference (±74%). If present, oil generally collects at the air-liquid interface and thus interferes with the alignment of the proteins and leads to a decrease in foam stability (Omowaye-Taiwo et al., 2015).

The mealworm powders showed superior foaming properties compared with those found in literature. For example, mealworm flours were reported to have a FC of 32% with a FS of 28% after 30 min (Zielińka et al., 2018). Kim et al. (2019) reported a FC of 130% and a FS of 78% (30 min) for water-soluble proteins extracted from defatted mealworm flours. However, as the results differ in method of determination and calculation, they are difficult to compare.

#### 3.2.3 Emulsion capacity and stability

To evaluate the emulsifying properties of the different mealworm powders, the EC and the ES were measured. The results, presented in **Figure 3(C-D)**, show that both parameters were significantly (p < 0.05) affected by fermentation. Without defatting, the EC decreased from 51% to 5% and 7% upon fermentation with the starters Bactoferm^®^ F-LC and *L. farciminis*, respectively, while the ES decreased from 74% to 0% and 18%. Reduction in emulsifying properties by fermentation has been observed earlier by Lampart-Szczapa et al. (2006) for lupin proteins. In this study, the lactic acid fermented lupin proteins were characterised by a lower hydrophobicity than non-modified lupin proteins and, as a consequence, by worse emulsifying properties. The emulsification properties of potato flours (Gong et al., 2019) and sorghum flour (Ogodo, Ugbogu, & Onyeagba, 2018), on the other hand, were improved by fermentation. Their soluble protein concentrations were increased during fermentation, which promotes oil droplet entrapment. Defatting significantly (p < 0.05) improved the EC of the fermented powders with 31% and 42% for FLC and Far, respectively, whereas the EC of the Control was unaffected. The ES was either improved (d-Control) or deteriorated (d-FLC and d-Far) by defatting.

The EC and ES of non-fermented mealworm powder are in line with those of other studies on mealworm flours (Lee et al., 2019; Zielińska et al., 2018). In addition, their emulsification properties were not significantly different from commercial egg albumen powder, indicating its potential as alternative source of protein emulsifier for food formulations.

#### 3.2.4 pH dependent protein solubility

**Figure 4** shows the protein solubility profiles of the mealworm powders in the pH range of 2 to 12. The protein solubility of the Control decreased in the pH range 2 to 4, showed a minimum solubility of 3% at pH 4, and gradually increased in the pH range 5 to 12. The highest solubility was found at pH 12 (77%). Using similar assay conditions, Bußler et al. (2016) and Zielińska et al. (2018) reported a maximum protein solubility of respectively 70% at pH 10 and 97% at pH 11. In contrast, less than 3% of the proteins were soluble near the isoelectric point (pH 4-5). Defatting of the Control did not lead to an increased yield of soluble proteins. On the contrary, protein solubility was significantly decreased at low (2-3) and high (8, 10-12) pH values. Regardless of the starter culture used, the fermentation process led to a drastic decrease in protein solubility and shifted the isoelectric point from pH 4 to pH 6. Similar results were obtained during the fermentation of sorghum by Elkhalifa, Schiffler, & Bernhardt (2005). In this study, lactic acid fermentation shifted the isoelectric point of sorghum proteins by two pH units and decreased the protein solubility due to the exposure of hydrophobic groups.

**Figure 4.**
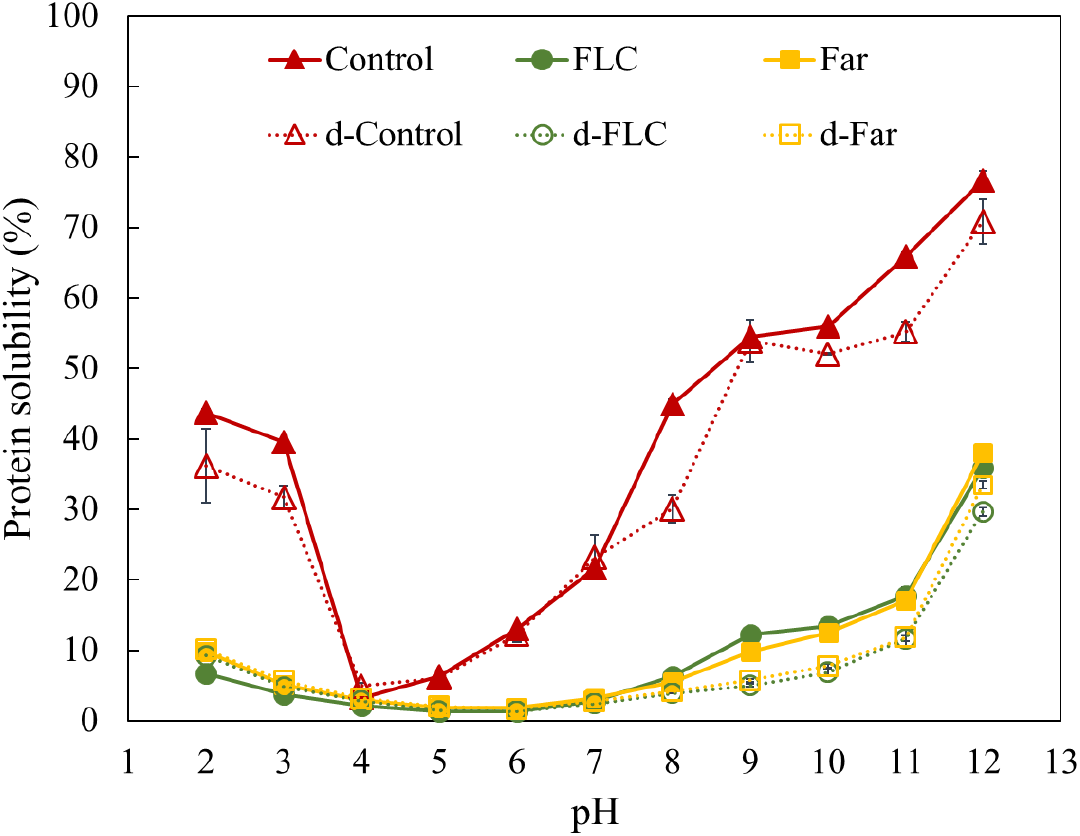
Protein solubility of the full fat (Control and the fermented samples FLC and FAR) and defatted (d-Control and the fermented samples d-FLC and d-Far) mealworm powders as a function of pH. Protein solubility [%] is presented to the total protein content analysed via Kjeldahl method.

Loss in solubility indicated denaturation or other structural changes of the mealworm proteins during processing. Both starter cultures tested in this study produced organic acids during fermentation as indicated by the pH reduction, which might have induced an irreversible coagulation of the proteins and thus a reduced solubility (Weng & Chen, 2010). Further, the fermentation process (including the blanching as pre-treatment) might have promoted aggregation and cross-linking of partially hydrolysed mealworms proteins, causing them to become insoluble (Paraman et al., 2007). In this context two molecular aspects need to be considered: on one side, due to partial hydrolysis occurring during the fermentation, the hydrophobic core of the proteins becomes exposed giving opportunity for aggregation of proteins based on non-covalent interactions. On the other side, the processing combined with fermentation generally leads to disulphide exchange between the exposed cysteine side chains, thus eventually promoting inter- and intramolecular covalent reactions. These two aspects promote the insolubility of the degradation products of the proteins.

#### 3.2.5 Quantification of free amino groups and protein molecular weight distribution

The amount of free amino groups found via the OPA assay in the protein extracts from the non-fermented powder d-Control (13.31 ± 2.19 mM/g soluble protein) was significantly (p < 0.05) lower than those of the fermented powders d-FLC (130.13 ± 2.06 mM/g soluble protein) and d-Far (144.49 ± 18.76 mM/g soluble protein). The high amount of free amino groups in the fermented samples may be attributed to proteolytic degradation of proteins during fermentation. Degradation of proteins resulting in an increase in free amino groups has been detected in many fermented products, such as yoghurt (Tavakoli et al., 2019), Suanyu (fermented fish, Wang et al., 2017) and mao-tofu (fermented soybean, Zhao & Zheng, 2009). The observed increase in free amino groups may point towards a corresponding increase in the free carboxyl groups resulting from the enzymatic degradation of the peptide bonds. Both these observations implicate the possibility of stronger ionic interactions, supporting the formation of non-covalent bonds during the aggregation discussed above, and thus making these molecular forms insoluble over a broader pH range.

SDS-PAGE analysis of the mealworm powders (Figure 5A, left figure) showed proteins with molecule weights between 10 and 250 kDa. For the non-fermented sample, the 55 kDa band was the most intensive in the lane. For the fermented powders, there was a band at 55 kDa, but between 70 and 100 kDa, there were also two clear bands. As can be seen in Figure 5A (right figure) and contrary to what can be expected, in relative terms fermented mealworm powders were more abundant in medium (25-70 kDa) and high (70-250 kDa) molecular weight proteins and less abundant in low-molecular weight proteins (0-25 kDa) than the non-fermented powders. This relative increase in medium and high molecular weight proteins by fermentation is probably to be explained by the fact that (1) during fermentation some of the low-molecular weight proteins are consumed by the starter cultures to ensure their growth (and as a result the relative abundances of medium/high molecular weight proteins increases) and/or that (2) the proteins with a molecular weight of 70 kDa or higher originate from the added starter culture cells. A further contribution to this observation may be related to macro protein structures that in the insects as such were not soluble, but become more accessible after the fermentation due to partial hydrolysis. Further in depth analysis is needed to characterize/identify the origin of these proteins and is envisaged in further studies. SDS-PAGE analysis of the soluble protein content of the non-fermented and fermented mealworm powders revealed that the fermentation process leads to a shift towards lower molecular weights of the proteins and that, depending on the pH, different protein fractions are soluble (Figure 5B-D). At pH 7, which was used for the evaluation of the emulsifying and foaming properties, the protein extract of the non-fermented mealworm powder was composed of 2.7% high molecular weight fraction, 72.1% medium molecular weight fraction and 9.8% low molecular weight fraction, whereas the protein fraction characterized by low molecular weights (0-15 kDa) were found to dominate the protein extracts of the fermented mealworm powders (57.4% and 80.9% for d-FLC and d-Far, respectively). As the molecular sizes of the soluble proteins decreased, it is clear that also the protein structure was modified by the fermentation process, which in turn leads to changes in techno-functional properties. Rahali et al. (2000) and Razali et al. (2015) reported that the surface hydrophobicity of the proteins is more important than the peptide length in emulsion and foaming properties. Most often, high surface hydrophobicity is needed to allow the formation of stable emulsions and foams. The hydrophilic/hydrophobic character of proteins is connected to their secondary, tertiary and quaternary structure and is caused by the amphiphilic character of amino acids and their accessibility in the polypeptide chain (Human et al., 2012). It could be hypothesized that the low molecular weight peptides produced by fermentation can migrate rapidly to the interface but that their hydrophilic/hydrophobic balance was insufficient for the stabilization of emulsions and foams.

**Figure 5.**
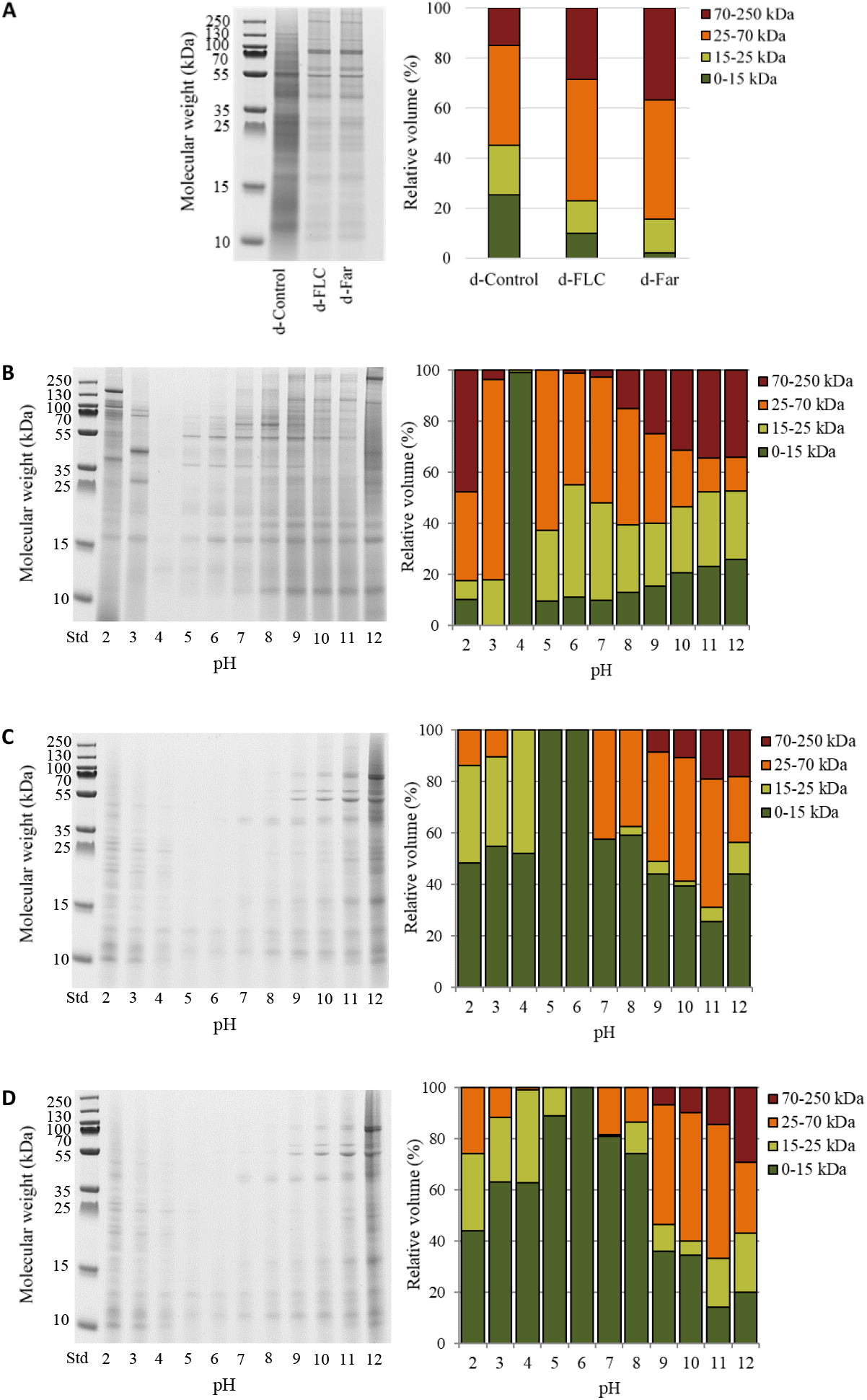
Gel electrophoresis (left) and relative composition (left) of mealworm powders (A) and soluble mealworm protein fractions (Control: regular mealworm paste (B), d-FLC: mealworm paste fermented with the starter Bactoferm^®^ F-LC (C), and d-Far: mealworm paste fermented with the starter Lactobacillus farciminis (D)) at different pH. Proteins are classified in four groups: Low molecular weight (0-15 kDa), medium molecular weight (15-25 kDa and 25-75 kDa) and high molecular weight (70-250 kDa).

## 4. Conclusions

Previous research has shown that fermentation of mealworm paste with lactic acid starter cultures results in a rapid pH reduction, as an indication of a successful fermentation process. In this study, the effect of fermentation on the functional properties was considered. When after the fermentation the flour is not defatted, WBC and OBC are decreased, the FC and FS are not affected and the EC and ES are reduced. When after the fermentation the flour is defatted, the same effects can be seen, with the exception that WBC and FS are (somewhat) improved compared to the non-fermented control. The differences in protein functionality between the control and the fermented powders may be ascribed to differences in molecular size of the proteins as indicated by the analysis of protein distribution and probably due to differences in hydrophilic/hydrophobic arrangement. It has to be concluded that fermentation with lactic acid starter cultures should not be pursued as a processing technology to obtain food ingredients based on mealworms for functional properties considered in this research. Nevertheless, this study confirmed that non-fermented mealworm powder has functional properties that are comparable to other protein sources. In addition, fermentation of mealworms and other insects will further be investigated as a technology for taste and shelf life improvement.

## Acknowledgements

This work was supported by internal KU Leuven funds (C32/16/024 e “Fermentation of edible insects”. The Alexander von Humboldt Foundation funded the author Sagu T.S (Ref 3.4 – CMR – 1164093 – GF-P).

## References

Ojokoh, A. O., Fayemi, O. E., Ocloo, F. C. K., & Nwokolo, F. I. (2015). Effect of fermentation on proximate composition, physicochemical and microbial characteristics of pearl millet (Pennisetum glaucum (L.) R. Br.) and Acha (Digitaria exilis (Kippist) Stapf) flour blends. Journal of Agricultural Biotechnology and Sustainable Development, 7(1), 1–8. https://doi.org/10.5897/jabsd2014.0236

Akpossan, R., Digbeu, Y., Koffi, M., Kouadio, J., Dué, E., & Kouamé, P. (2015). Protein Fractions and Functional Properties of Dried Imbrasia oyemensis Larvae Full-Fat and Defatted Flours. International Journal of Biochemistry Research & Review, 5(2), 116—126. https://doi.org/10.9734/ijbcrr/2015/12178

Borremans, A., Crauwels, S., Vandeweyer, D., Smets, R., Verreth, C., Van Der Borght, M., Lievens, B., & Van Campenhout, L. (2019). Comparison of six commercial meat starter cultures for the fermentation of yellow mealworm (Tenebrio molitor) paste. Microorganisms, 7(11). https://doi.org/10.3390/microorganisms7110540

Borremans, A., Lenaerts, S., Crauwels, S., Lievens, B., & Van Campenhout, L. (2018). Marination and fermentation of yellow mealworm larvae (Tenebrio molitor). Food Control, 92, 47–52. https://doi.org/10.1016/j.foodcont.2018.04.036

Broyard, C., & Gaucheron, F. (2015). Modifications of structures and functions of caseins: a scientific and technological challenge. Dairy Science and Technology, 95(6), 831–862. https://doi.org/10.1007/s13594-015-0220-y

Bußler, S., Rumpold, B. A., Jander, E., Rawel, H. M., & Schlüter, O. K. (2016). Recovery and techno-functionality of flours and proteins from two edible insect species: Meal worm (Tenebrio molitor) and black soldier fly (Hermetia illucens) larvae. Heliyon, 2(12). https://doi.org/10.1016/j.heliyon.2016.e00218

Çabuk, B., Nosworthy, M. G., Stone, A. K., Korber, D. R., Tanaka, T., House, J. D., & Nickerson, M. T. (2018). Effect of fermentation on the protein digestibility and levels of non-nutritive compounds of pea protein concentrate. Food Technology and Biotechnology, 56(2), 257–264. https://doi.org/10.17113/ftb.56.02.18.5450

De Smet, J., Lenaerts, S., Borremans, A., Scholliers, J., Van Der Borght, M., & Van Campenhout, L. (2019). Stability assessment and laboratory scale fermentation of pastes produced on a pilot scale from mealworms (Tenebrio molitor). Lwt, 102(November 2018), 113–121. https://doi.org/10.1016/j.lwt.2018.12.017

Gong, S., Xie, F., Lan, X., Zhang, W., Gu, X., & Wang, Z. (2019). Effects of Fermentation on Compositions, Color, and Functional Properties of Gelatinized Potato Flours. Journal of Food Science, 85, 57–64. https://doi.org/10.1111/1750-3841.14837

Gravel, A., & Doyen, A. (2020). The use of edible insect proteins in food: Challenges and issues related to their functional properties. Innovative Food Science and Emerging Technologies, 59(October 2019), 102272. https://doi.org/10.1016/j.ifset.2019.102272

Human, C., Aquaporins, P., & Nordén, K. (2012). From Sequence to Structure. 1–47.

Kim, H. W., Setyabrata, D., Lee, Y. J., Jones, O. G., & Kim, Y. H. B. (2016). Pre-treated mealworm larvae and silkworm pupae as a novel protein ingredient in emulsion sausages. Innovative Food Science and Emerging Technologies, 38, 116–123. https://doi.org/10.1016/j.ifset.2016.09.023

Klupsaite, D., Juodeikiene, G., Zadeike, D., Bartkiene, E., Maknickiene, Z., & Liutkute, G. (2017). The influence of lactic acid fermentation on functional properties of narrow-leaved lupine protein as functional additive for higher value wheat bread. LWT - Food Science and Technology, 75, 180–186. https://doi.org/10.1016/j.lwt.2016.08.058

Lampart-Szczapa, E., Konieczny, P., Nogala-Kałucka, M., Walczak, S., Kossowska, I., & Malinowska, M. (2006). Some functional properties of lupin proteins modified by lactic fermentation and extrusion. Food Chemistry, 96(2), 290–296. https://doi.org/10.1016/j.foodchem.2005.02.031

Lee, H., Kim, J., Ji, D., & Lee, C. (2019). Effects of Heating Time and Temperature on Functional Properties of Proteins of Yellow Mealworm Larvae (Tenebrio molitor L.). Food Science of Animal Resources, 39(2), 296–308. https://doi.org/10.5851/kosfa.2019.e24

Lenaerts, S., Van Der Borght, M., Callens, A., & Van Campenhout, L. (2018). Suitability of microwave drying for mealworms (Tenebrio molitor) as alternative to freeze drying: Impact on nutritional quality and colour. Food Chemistry, 254(February), 129–136. https://doi.org/10.1016/j.foodchem.2018.02.006

Malomo, S. A., He, R., & Aluko, R. E. (2014). Structural and functional properties of hemp seed protein products. Journal of Food Science, 79(8). https://doi.org/10.1111/1750-3841.12537

Nkhata, S. G., Ayua, E., Kamau, E. H., & Shingiro, J. B. (2018). Fermentation and germination improve nutritional value of cereals and legumes through activation of endogenous enzymes. Food Science and Nutrition, 6(8), 2446–2458. https://doi.org/10.1002/fsn3.846

Ogodo, A. C., Ugbogu, O. C., & Onyeagba, R. A. (2018). Variations in the functional properties of soybean flour fermented with lactic acid bacteria. International Journal of Biology and Biomedical Engineering, 12(January), 1–6. https://doi.org/10.4172/2471-9315.1000141

Ogodo, A. C., Ugbogu, O. C., Onyeagba, R. A., & Okereke, H. C. (2018). In-vitro starch and protein digestibility and proximate composition of soybean flour fermented with lactic acid bacteria (LAB) consortia. Agriculture and Natural Resources, 52(5), 503–509. https://doi.org/10.1016/j.anres.2018.10.001

Omowaye-Taiwo, O. A., Fagbemi, T. N., Ogunbusola, E. M., & Badejo, A. A. (2015). Effect of germination and fermentation on the proximate composition and functional properties of full-fat and defatted cucumeropsis mannii seed flours. Journal of Food Science and Technology, 52(8), 5257–5263. https://doi.org/10.1007/s13197-014-1569-2

Paraman, I., Hettiarachchy, N. S., Schaefer, C., & Beck, M. I. (2007). Hydrophobicity, solubility, and emulsifying properties of enzyme-modified rice endosperm protein. Cereal Chemistry, 84(4), 343–349. https://doi.org/10.1094/CCHEM-84-4-0343

Rahali, V., Chobert, J. M., Haertlé, T., & Guéguen, J. (2000). Emulsification of chemical and enzymatic hydrolysates of β-lactoglobulin: Characterization of the peptides adsorbed at the interface. Nahrung - Food, 44(2), 89–95. https://doi.org/10.1002/(sici)1521-3803(20000301)44:2<89::aid-food89>3.3.co;2-l

Razali, A. N., Amin, A. M., & Sarbon, N. M. (2015). Antioxidant activity and functional properties of fractionated cobia skin gelatin hydrolysate at different molecular weight. International Food Research Journal, 22(2), 651–660.

Rumpold, B. A., & Schlüter, O. K. (2013). Potential and challenges of insects as an innovative source for food and feed production. Innovative Food Science and Emerging Technologies, 17, 1–11. https://doi.org/10.1016/j.ifset.2012.11.005

Tavakoli, M., Habibi Najafi, M. B., & Mohebbi, M. (2019). Effect of the milk fat content and starter culture selection on proteolysis and antioxidant activity of probiotic yogurt. Heliyon, 5(2). https://doi.org/10.1016/j.heliyon.2019.e01204

Van Huis, A. (2016). Edible insects are the future? Proceedings of the Nutrition Society, 75(3), 294–305. https://doi.org/10.1017/S0029665116000069

Wang, W., Xia, W., Gao, P., Xu, Y., & Jiang, Q. (2017). Proteolysis during fermentation of Suanyu as a traditional fermented fish product of China. International Journal of Food Properties, 20(1), S166–S176. https://doi.org/10.1080/10942912.2017.1293089

Yi, L., Lakemond, C. M. M., Sagis, L. M. C., Eisner-Schadler, V., Huis, A. Van, & Boekel, M. A. J. S. V. (2013). Extraction and characterisation of protein fractions from five insect species. Food Chemistry, 141(4), 3341–3348. https://doi.org/10.1016/j.foodchem.2013.05.115

Zhao, Xinhuai, & Zheng, X. (2009). A primary study on texture modification and proteolysis of mao-tofu during fermentation. African Journal of Biotechnology, 8(10), 2294–2300. https://doi.org/10.5897/AJB2009.000-9293

Zhao, Xue, Vázquez-Gutiérrez, J. L., Johansson, D. P., Landberg, R., & Langton, M. (2016). Yellow mealworm protein for food purposes - Extraction and functional properties. PLoS ONE, 11(2), 1–17. https://doi.org/10.1371/journal.pone.0147791

Zielińska, E., Karaś, M., & Baraniak, B. (2018). Comparison of functional properties of edible insects and protein preparations thereof. LWT - Food Science and Technology, 91(October 2017), 168–174. https://doi.org/10.1016/j.lwt.2018.01.058

