## Supplementary material for "Functional properties of powders produced from either or not fermented mealworm (*Tenebrio molitor*) paste": Table S1 Supporting Information

**Table 1 (Supporting information).** Initial protein concentration, quantity of protein loaded on the gel and relative and absolute abundances of the soluble proteins with molecular weight 0-15 kDA of non-fermented (d-Control) and fermented (d-FAR and d-FLC) mealworm powders.

| Well No | 1 | 2 | 3 | 4 | 5 | 6 | 7 | 8 | 9 | 10 | 11 | 12 |
|---|---|---|---|---|---|---|---|---|---|---|---|---|
| Sample | Std | pH 2 | pH 3 | pH 4 | pH 5 | pH 6 | pH 7 | pH 8 | pH 9 | pH 10 | pH 11 | pH 12 |
| **D-Control** | | | | | | | | | | | | |
| Initial protein concentration (µg/ml) | / | 2097.1 | 2028.6 | 314.3 | 387.5 | 780.4 | 1480.1 | 1899.1 | 3448.9 | 3317.3 | 3479.8 | 4494.7 |
| Quantity of protein loaded (µg) | / | 6.29 | 6.09 | 0.94 | 1.16 | 2.34 | 4.44 | 5.7 | 10.35 | 9.95 | 10.44 | 13.48 |
| MW 0-15 kDA (relative) | | 14 | 0 | 96.5 | 8.14 | 15.11 | 13.95 | 16.28 | 18.60 | 24.41 | 26.74 | 29.07 |
| MW 0-15 kDA (absolute) | | 0.88 | 0 | 0.97 | 0.09 | 0.35 | 0.62 | 0.93 | 1.92 | 2.43 | 2.79 | 3.91 |
| **d-FLC** | | | | | | | | | | | | |
| Initial protein concentration (µg/ml) | / | 575.3 | 267.9 | 162.4 | 81.7 | 78.7 | 142.5 | 225.3 | 284.3 | 387.8 | 653.4 | 1677.4 |
| Quantity of protein loaded (µg) | / | 1.78 | 0.89 | 0.54 | 0.27 | 0.26 | 0.48 | 0.75 | 0.95 | 1.29 | 2.18 | 5.59 |
| MW 0-15 kDA (relative) | | 52.38 | 57.14 | 54.76 | 100 | 100 | 60.71 | 61.90 | 47.62 | 42.86 | 29.76 | 47.62 |
| MW 0-15 kDA (absolute) | | 0.93 | 0.51 | 0.30 | 0.27 | 0.26 | 0.29 | 0.46 | 0.45 | 0.55 | 0.64 | 2.66 |
| **d-Far** | | | | | | | | | | | | |
| Initial protein concentration (µg/ml) | / | 533.4 | 323.3 | 181.5 | 116.2 | 96.7 | 156.2 | 233.6 | 320.6 | 445.2 | 683 | 1884.1 |
| Quantity of protein loaded (µg) | / | 1.92 | 1.08 | 0.61 | 0.39 | 0.32 | 0.52 | 0.78 | 1.07 | 1.48 | 2.28 | 6.28 |
| MW 0-15 kDA (relative) | | 47.06 | 63.53 | 63.53 | 89.41 | 100 | 81.18 | 75.29 | 40 | 37.65 | 16.47 | 23.3 |
| MW 0-15 kDA (absolute) | | 0.61 | 0.69 | 0.39 | 0.35 | 0.32 | 0.42 | 0.59 | 0.43 | 0.56 | 0.38 | 1.46 |
